# Glydentify: An explainable deep learning platform for glycosyltransferase donor substrate prediction

**DOI:** 10.64898/2026.03.13.711695

**Authors:** Ruili Fang, Lan Na, Charles J. Corulli, Pradeep K. Prabhakar, Steven J Berardinelli, Aarya Venkat, Anup Prasad, Rezwan Mahmud, Kelley W. Moremen, Breeanna R. Urbanowicz, Fei Dou, Natarajan Kannan

## Abstract

Glycosyltransferases (GTs) are a large family of enzymes that catalyze glycosidic linkages formation between chemically diverse donor and acceptor molecules to regulate diverse cellular processes across all domains of life. Despite their importance, the activated sugar donors (donor substrates) used by most GTs remain unidentified, limiting our understanding of GT functions. To address this challenge, we developed Glydentify, a deep learning framework that predicts donor usage across GT-A and GT-B fold glycosyltransferases. Trained on large-scale UniProt annotations, Glydentify integrates protein sequence embeddings learned from protein language models with chemical features derived from molecular encoders trained on extensive chemical datasets. The resulting models achieve high predictive performance, with precision–recall AUCs (PR-AUC) of 0.86 for GT-A and 0.91 for GT-B, surpassing general enzyme–substrate predictors while requiring minimal manual curation. We employed Glydentify to predict the donor specificity of uncharacterized plant GTs and experimentally tested the predictions using in vitro biochemical assays. Furthermore, we demonstrate that the model utilizes a combination of evolutionary, structural, and biochemical features to predict donor specificity through residue attention score analysis. Together, these results establish Glydentify as a robust, explainable framework for decoding donor-glycosyltransferase relationships and highlight its potential as a broadly applicable framework for modeling enzyme classes that act on chemically diverse substrates.

## Introduction

Glycosylation is a ubiquitous and fundamental mechanism for regulating diverse biological processes across all domains of life ^1,2^. The diverse biological roles of glycosylation are directly linked to the remarkable chemical diversity of glycans^3,4^. Glycosylation is catalyzed by glycosyltransferases (GTs), a large family of enzymes that transfer sugar moieties from activated nucleotide-sugar donors to a wide range of acceptor molecules, including proteins, lipids, small molecules, and other carbohydrates. Over evolutionary time, GTs have diversified to utilize a broad spectrum of chemically distinct nucleotide sugars, shown in **Error! Reference source not found.**. Two major structural folds, GT-A and GT-B, dominate the known GT landscape and rely on nucleotide sugars as donors^5^. Despite their central role in glycan biosynthesis, donor usage remains unknown for the vast majority of putative GTs. The CAZy database catalogs more than 700,000 GT entries from diverse organisms (to this date), grouped into 139 families. However, only a small fraction of these families have been experimentally characterized with respect to their donor substrates. An incomplete understanding of GT sugar donor specificity presents a major challenge in engineering GTs for biotechnological applications, such as antibody engineering, vaccine development, and modulation of natural-product pharmacokinetics^1^.

Biochemical assays for donor usage are costly, low-throughput, and generally limited to a narrow panel of candidate sugars, making comprehensive mapping impractical. Moreover, donor specificity cannot be inferred reliably from sequence or structure alone. Sugar donor specificity is often governed by spatially dispersed, co-evolving residues and epistatic interactions, which cannot be inferred through simple sequence patterns or structural comparisons^5^. For instance, enzymes adopting entirely distinct structural scaffolds, such as cellulose synthase from family GT2 (GT-A fold)^6^ and glycogen synthase, from family GT3 or GT5 (GT-B fold)^7^, can both bind UDP-glucose; in contrast, in the ABO blood group transferase, only four amino acid substitutions convert specificity from UDP–GalNAc to UDP–Gal ^8^. This challenge is further compounded by the high physicochemical similarity of nucleotide-sugar donors, which share a conserved nucleotide moiety (e.g., UDP) and differ only by subtle stereochemical changes at one carbon in the hexose ring(**Figure 1A**). This structural similarity presents a significant challenge for predictive modeling, as enzymes often exhibit strict stereochemical stringency, where even minor stereochemical variations (e.g., the C4-epimerization between glucose and galactose) can completely abolish substrate recognition. Such cryptic determinants highlight why homology-based annotations are inadequate for predicting sugar donor specificity. Thus, there is a need for specialized tools to advance glycoenzyme function prediction.

**Figure 1.**
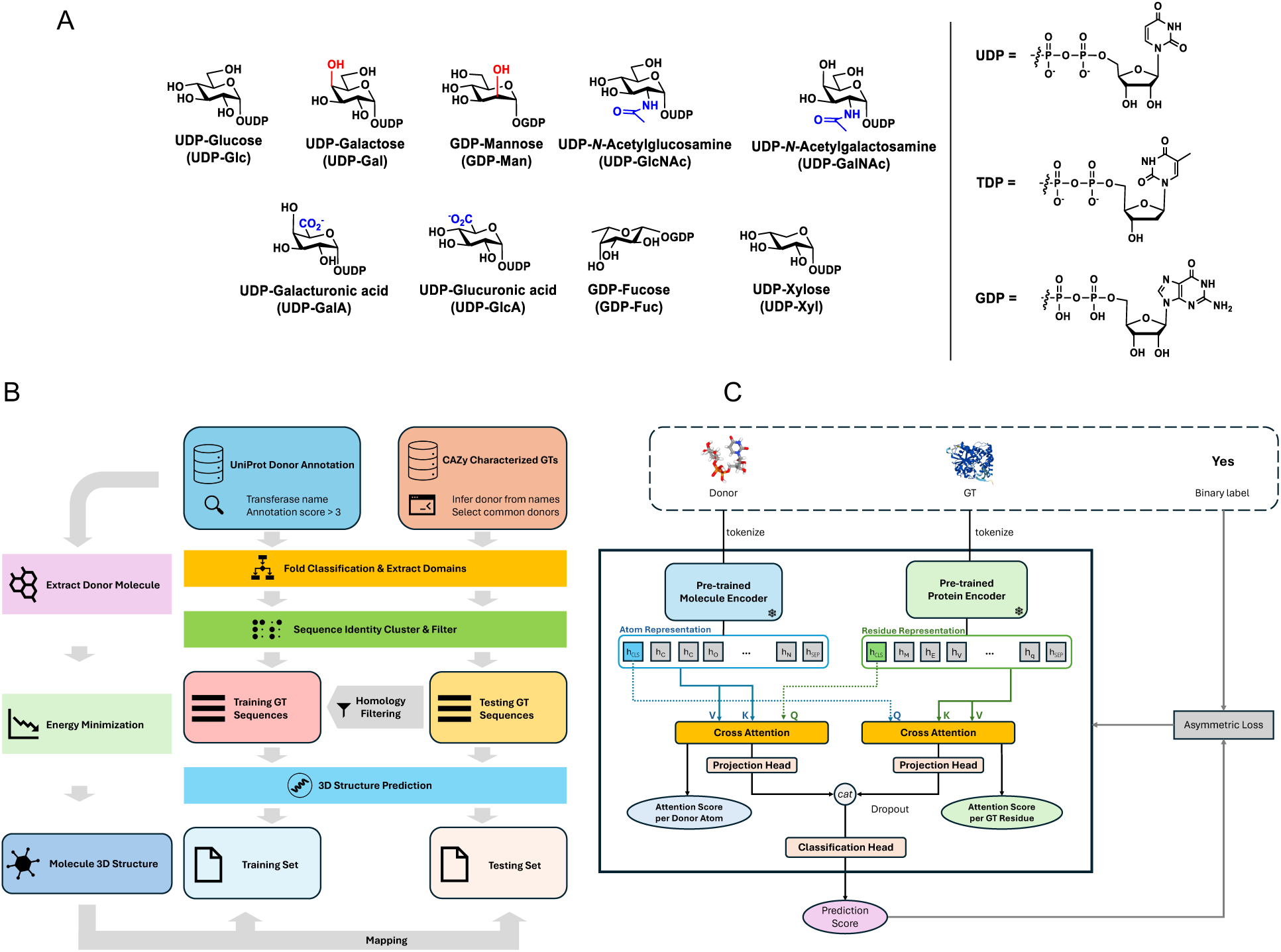
Donor diversity and overview of Glydentify model architecture and training. **A.** Chemical diversity of the sugar donor. Chemical structure of sugar donor used in the current prediction was shown. **B.** A full schematic of our data curation pipelines. **C.** The architecture of the Glydentify model classifies GT-Donor pairs as positive or negative interactions.

Previous computational approaches for predicting sugar donor specificity in GTs largely relied on statistical models and handcrafted features, restricting their application to specific families or fold-types. Yang et al.^9^ introduced GT-Predict to model Arabidopsis GT1 enzymes with decision trees that combined physicochemical descriptors of both donors and acceptors. Taujale et al.^10^ predicted donor classes within GT-A enzymes using residue-level physicochemical descriptors derived from curated sequence alignments. Moreover, many supervised learning approaches have framed such problems as single-label classification tasks, assuming one donor per enzyme. For instance, Hennen et al.^11^ trained GT-B classifiers on curated data using sequence properties and AF2-based structural features, but assumed that one GT could only utilize one donor sugar. This overlooks the biological reality that some GTs exhibit dual or promiscuous donor specificity. While demonstrating feasibility, these models remain constrained by the reliance on handcrafted input features, limited scope, and single-label formulations.

Conversely, the advent of deep learning and protein language models has revolutionized protein function prediction tasks by learning biological representations directly from exiting data in an unsupervised way, eliminating the need for manual feature curation. Initial efforts in the glycoenzyme space, such as DeepGlycanSite^12^, GlyNet^13^, SweetNet^14^, and CandyCrunch, leveraged Graph Neural Networks (GNNs) to capture the subtle sequence–structure interactions governing glycan–protein interactions. Furthermore, protein-language models like ESM-2^15^ utilize self-supervised methods on millions of publicly available sequences to extract context-aware representations for downstream tasks. Such pre-trained protein language models have been successfully adapted by enzyme-substrate predictors such as ESP^16^ and EZspecificity^17^. More recent protein-language models, such as ESM-^18^ and SaProt^19^, further extend the paradigm by incorporating explicit structural information during pretraining.

Despite these recent advancements in protein function prediction, representation learning approaches have not yet been fully harnessed for GT donor substrate prediction. General frameworks, such as ESP^16^, broadly cover enzyme substrates, and the enzyme specificity predictor EZspecificit^17^ has been tested on some glycosyltransferase acceptor substrates. However, these models lack the resolution to predict GT donor specificity at scale, primarily due to the limited GT data available in the general enzyme databases used for training. Additionally, these models lack interpretability, limiting their applications in investigating the relationships connecting sequence, structure, and function in GTs. To bridge this gap, we develop Glydentify, an explainable evotuned deep learning framework trained on GT-A and GT-B fold families from diverse organisms. The model leverages pre-trained protein and donor embeddings in an end-to-end framework, utilizing bi-directional cross-attention to support multi-label donor prediction across different GT families and fold classes, while learning from existing data in an unsupervised manner. Unlike earlier approaches, Glydentify does not rely on handcrafted descriptors or sequence alignments; instead, it learns directly from the intrinsic structure and context of GTs and their donor substrates. The framework estimates the probability of each donor sugar independently, enabling the identification of enzymes with multiple donor specificities. Trained on large-scale UniProt annotations and evaluated against CAZy entries, Glydentify achieves state-of-the-art predictive performance while offering interpretability in identifying protein residues that contribute to donor prediction. We demonstrate the application of Glydentify in predicting and experimentally evaluating donor substrates in a subset of GT-A and GT-B plant transferases not used in training. By establishing a robust and scalable baseline for donor prediction, Glydentify offers a practical tool for functional annotation of GTs and a tool for guiding experimental discovery in glycobiology.

## Results

### Glydentify: Attention-guided weighting of GT and donor fused representation

To address the challenges of predicting GT donor specificity at scale, we developed Glydentify, a structure-aware deep learning framework that integrates protein and donor information. While explicitly modeling enzyme-substrate complex structures can offer the most rigorous basis for defining specificity^17^, the scarcity of experimentally verified complex structures makes it difficult to train a deep learning model. Moreover, computational surrogates could introduce noise and eventually compromise a model’s performance. Generative models like AlphaFold 3^20^ frequently exhibit stereochemical inaccuracies^21,22^, whereas molecular docking optimizes for ground-state binding stability rather than the transition-state geometry required for turnover. In contrast, Glydentify captures the high-dimensional chemical logic of donor recognition without relying on these noisy structural priors. Two separate predictors were trained for GT-A and GT-B enzymes. Each predictor has the same architecture as shown in **Figure 1B**. A pretrained protein language model encodes each GTs information at the residue level. In parallel, a pretrained molecule encoder represents donor sugars at atomic resolution. To integrate these modalities, we employ a bi-directional cross-attention module that enables residues and donor atoms to exchange information, assigning attention weights to prioritize informative features for donor recognition. A final classifier produces probability scores for each GT-donor pair, which, when concatenated across donors, yields a multi-label output vector for each GT. This design allows Glydentify to learn directly from protein sequences and donor molecules, bypassing the need for handcrafted features or sequence alignments.

We systematically evaluated different protein language models to understand how sequence-versus structure-aware representations influence donor sugar prediction. We hypothesized that protein language models trained under different objectives capture distinct biochemical features of glycosyltransferases, leading to variations in how donor-recognition patterns are represented and ultimately predicted.

Specifically, ESM-2 was trained on large-scale natural protein sequences to capture evolutionary co-variation, ESM-C was contrastively trained to emphasize structural and physicochemical consistency, and SaProt incorporated AlphaFold-derived embeddings to encode explicit three-dimensional geometric information. To further enhance structural awareness across all representations, we generated AlphaFold3^20^ predictions for all training and test sequences. These were used to enrich the SaProt sequence embeddings with structural context during the representation learning process. Donor molecules were encoded from SMILES strings into 3D conformers using RDKit and UniMol V2, ensuring that the chemical diversity of nucleotide sugars was consistently represented.

We trained Glydentify on a large set of UniProt entries annotated with donor usage (**Figure 1C**). To reduce label noise, only entries with high annotation scores were retained, and each entry was categorized as GT-A or GT-B based on HMM profiles. To prevent redundancy, we clustered sequences within each donor type, and then selected representatives at 95–100% identity thresholds depending on donor sample size. For evaluation, we constructed a benchmark test set from CAZy by extracting experimentally verified GT–donor pairs, clustering at 90% identity, and removing any training homologs with > 90% sequence identity to any test sequence. This split ensures that performance is assessed on clean labels and different sequence similarities.

For each enzyme, the model outputs an independent probability for every donor in the vocabulary. Training was performed under a closed-world assumption, where unannotated donors are treated as negatives. Although this may introduce occasional false negatives, the effect is mitigated by the large ratio of true negatives to positives, providing a consistent and reproducible setup across donor classes.

With this framework and dataset in place, we next evaluated Glydentify’s global performance on the CAZy benchmark, comparing it to baseline sequence-only models and to a general enzyme–substrate predictor.

### Glydentify outperforms existing machine learning models for donor substrate prediction

We assessed the performance of Glydentify by comparing its results with existing general-purpose models. To further validate the effectiveness of the architecture, we also tested a standard classification pipeline consisting of a pre-trained protein encoder backbone with an MLP head trained on our own data. We found that the general enzyme substrate predictors ESP^16^ and EZspecificity^17^ perform only slightly above random, reflecting the extreme sparsity of glycosyltransferase-donor data in its training data (**Figure 2A**). This limitation motivated our curation of a focused GT donor dataset. A simple pre-trained encoder with a multi-label classification head trained on our UniProt-derived GT sequences already delivers strong gains (PR-AUC ≈ 0.70, ROC-AUC ≈ 0.80, MCC ≈ 0.63), indicating that sequence information alone carries substantial signal. When integrating the rich-context embedding from the pre-trained protein encoder and small molecule encoder via cross-attention, the performance can be further improved. Although the overall performances across different encoder choices are similar, Glydentify with SaProt and UnimolV2 achieves the best performance across all metrics (PR-AUC: 0.82–0.90, ROC-AUC: 0.94–0.97, MCC: 0.71–0.81). Furthermore, these performance gains are consistent on both GT-A and GT-B fold enzymes, demonstrating the models ability to resolve donor specificity under heavy class imbalance.

**Figure 2.**
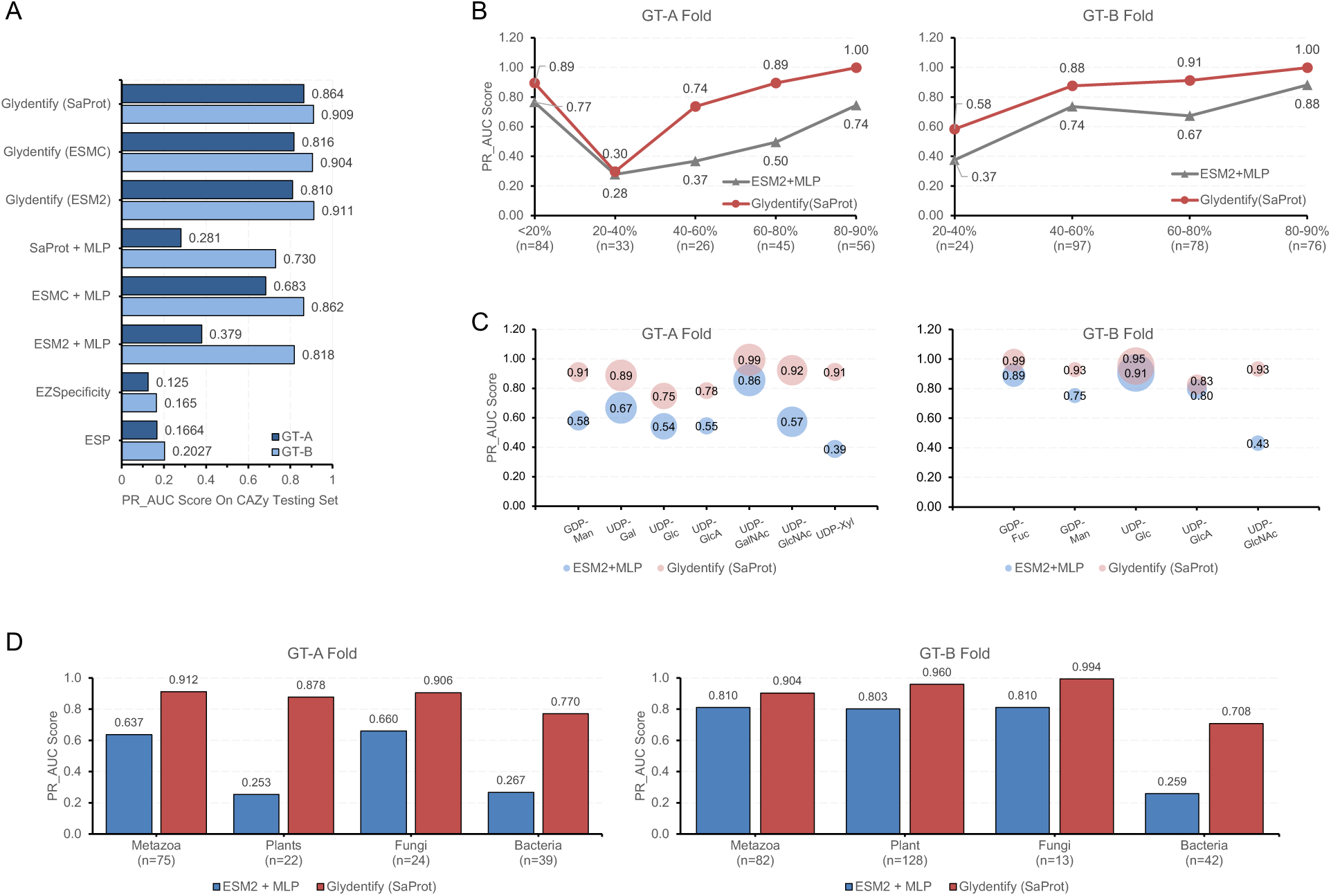
Evaluation of Glydentify performance. **A**. Overall Performance comparing with ESP, EZspecificity, predicting with only pre-trained protein language models and our designs with different pre-trained protein language models. All pre-trained encoders are frozen and only the shallow MLP probe or fusion head is trained. **B**. Performance as a function of sequence similarity between each test enzyme and its closest training sequence (Methods), binned as <20%, 20–40%, 40–60%, 60–80%, and 80–90%. Bars report PR-AUC; the dashed line (right y-axis) shows the number of GTs per bin. **C.** Donor-wise performance for GT-A (left) and GT-B (right). Bars show PR-AUC for the models indicated in the legend; the dashed line (right y-axis) gives the number of positive sequences available for each donor in the evaluation set. **D.** Performance by taxonomic group for GT-A (left) and GT-B (right). Bars report PR-AUC; the dashed line (right y-axis) indicates the number of GTs in each kingdom. Donor/taxon/identity categories with fewer than 10 test positives are omitted because PR–AUC is unreliable at very limited sample sizes.

Having established that models trained on a GT-specific training set substantially outperform boths general enzyme-substrate predictors, we next examine how performance varies across biologically and data-driven subgroups. This is critical since global averages can conceal systematic weakness on under-represented donors, rare taxa or remote homologs. Therefore, we stratified the CAZy test sets along three different categories: sequence novelty (percent identity to the nearest training example, binned at 0–20%, 20–40%, 40–60%, 60-80%, 80-90%), donor sugar type, and taxonomic origin (kingdom). For each subgroup and fold (GT-A/GT-B), we report PR-AUC as the primary metric. To aid interpretation, each panel overlays the number of (positive) samples of that subgroup in the training set and testing set. Subgroups with fewer than 10 test positives are excluded because PR-AUC becomes unstable at very small values. Overall, our data showed that *Glydentify* outperforms the protein-encoder-only baselines across all donor classes that meet the inclusion criterion. Across both folds, PR-AUC increases with sequence identity, and Glydentify (red) outperforms the ESM-2–only baseline (blue) in every bin that meets the inclusion criterion (**Figure 2B**). Gains are largest in the low–identity regime, which is precisely where the distribution shift is strongest. The monotonic trend with identity and consistent gains in the 20–60% bins indicate that cross–attending SaProt embeddings with UniMol donor geometry mitigate generalization gaps to remote homologs, while maintaining near–ceiling performance when close sequence neighbors are present. For performance across different donor sugars, while the model achieves high PR-AUC for well-represented donor types (**Figure 2C**), we observed a distinct performance drop-off for rare donors (N<50) within the training set (**Supplementary Table 2**), similar to the limitations observed in general enzyme-substrate models. To systematically define this data dependency, we performed a few-shot fine-tuning analysis (**Supplementary Text 1**). Both the minor-donor evaluation and the few-shot experiments converge on a similar finding: successful generalization requires a critical threshold of annotated examples (approx. N>50), below which the structural logic of donor specificity cannot be robustly inferred from sequence alone. As for the performance on different organisms, across both folds, Glydentify consistently exceeds the ESM-2–only baseline (blue) in the major, well–represented clades (**Figure 2D**). Gains are largest where we have ample supervision or a clear donor–sequence signal. Performance on *Metazoan* and plant (*Viridiplantae)* GTs reaches near–ceiling PR–AUC, with substantial improvements for GT–B fold enzymes (e.g., *Metazoa* + ∼0.19 PR–AUC, *Viridiplantae* +∼0.08). For proteins that adopt a GT-A fold, predictions on fungal GTs showed a pronounced improvement (+∼0.25), while predictions on bacterial GTs (*Pseudomonadota)* improved modestly. Mixed outcomes appear in very small bacterial/viral groups (e.g., *Bacillota*, *Mycoplasmatota*); their PR–AUCs have high variance due to the small number of positive training samples in those categories, so we did not overinterpret those fluctuations. Overall, the kingdom-stratified analysis indicates that explicit protein–donor interaction modeling transfers well across eukaryotic taxa and moderately across bacteria, with variability in performance likely reflecting biological divergence among kingdoms rather than differences in dataset size.

### Evaluation on independent datasets demonstrates Glydentify’s generalizability across diverse species and families

We evaluated Glydentify’s performance on two independent datasets spanning distinct GT folds and GT families to assess its generalizability beyond the training distribution (**Table 1**). For GT-A fold proteins, we curated a literature-based set of experimentally validated GTs with known donor specificities. For enzymes that adopt a GT-B fold, we focused on *Spirodela polyrhiza* and *Spirodela intermedia* enzymes from the GT-47 family. The selection of plant GT47 enzymes for this study was due to their broad functional diversity. More specifically, plant GT47 members are known to have the capacity to utilize a diverse array of activated nucleotide sugar donor substrates, including UDP-β-L-Araf, UDP-β-L-Arap, UDP-α-D-Xyl, UDP-α-D-Gal, and UDP-α-D-GalA. Furthermore, they are known or hypothesized to be involved in the synthesis of all major classes of plant cell wall matrix polysaccharides; however, many of their donor and acceptor specificities remain unknown, making functional characterization efforts difficult^23^. To validate the donor prediction accuracy of the Glydentify model within this family, candidate enzymes were heterologously expressed in HEK 293 cells and purified. Nucleotide sugar donor specificity was subsequently determined by quantifying the release of UDP in the absence of an acceptor, a byproduct of GT activity after a sugar from the activated donor is transferred to water^24^, providing an experimental benchmark for the model’s predictive performance. Enzymes that exhibited no hydrolytic activity in the absence of acceptor were assayed with an appropriate acceptor substrate to identify donor specificity.

**Table 1.** Independent validation of Glydentify predictions on experimentally verified and new GTs. The table lists representative GT-A and GT-B enzymes from independent datasets, along with their known donor sugars, sequence similarity to the training set, and predicted donor-specificity scores.

| FOLD | NAME OR UNIPROT ID | ORGANISM | VALIDATED DONOR SUGAR | MAX SEQ. IDENT. (%) | MODEL'S PREDICTION (%) |
| --- | --- | --- | --- | --- | --- |
| GT-A | J9VFZ3 | Cryptococcus neoformans | GDP-Man | 26.5 | GDP-Man (90.9); UDP-Glc (54.4) |
|  | Q72K30 | Thermus thermophilus HB27 | GDP-Man | 49.4 | GDP-Man (79.0) |
|  | P53059 | Saccharomyces cerevisiae S288c | GDP-Man | 48.1 | GDP-Man (99.6) |
| GT-B | Sp146-A | *Spirodela polyrhiza* | UDP-Gal | 64 | UDP-Gal (98.4) |
|  | Sp197-A | *Spirodela polyrhiza* | UDP-Gal | 45 | UDP-Gal (99.4) |
|  | Sp291-D | *Spirodela polyrhiza* | UDP-Xyl | 63.2 | UDP-GlcA (77.0); UDP-Xyl (51.3) |
|  | Sp415-C | *Spirodela polyrhiza* | UDP-Xyl | 50 | UDP-Xyl (93.3) |
|  | Sp124-C | *Spirodela polyrhiza* | UDP-Xyl | 44.3 | UDP-Xyl (84.3) |
|  | Sp124-E | *Spirodela polyrhiza* | UDP-Xyl | 37.6 | UDP-Xyl (73.4); UDP-GlcA (52.1) |
|  | Si728-C | *Spirodela intermedia* | UDP-Xyl | 48.8 | UDP-Xyl (93.3); UDP-GlcA (61.9) |

Despite low sequence similarity to the current training set (**Table 1**), Glydentify accurately predicted donor sugar identity across the newly characterized enzymes. Notably, the recently described CAZy family GT-139 enzyme Cgm1 (Uniprot ID: J9VFZ3), a mannosyltransferase involved in the biosynthesis of capsular glucuronoxylomannogalactan in *Cryptococcus neoformans*^25^, was correctly assigned GDP-Man as its preferred donor with 90% prediction score (**Figure 3A**). This result demonstrates Glydentify’s robust ability to generalize to even newly added CAZy families.

**Figure 3.**
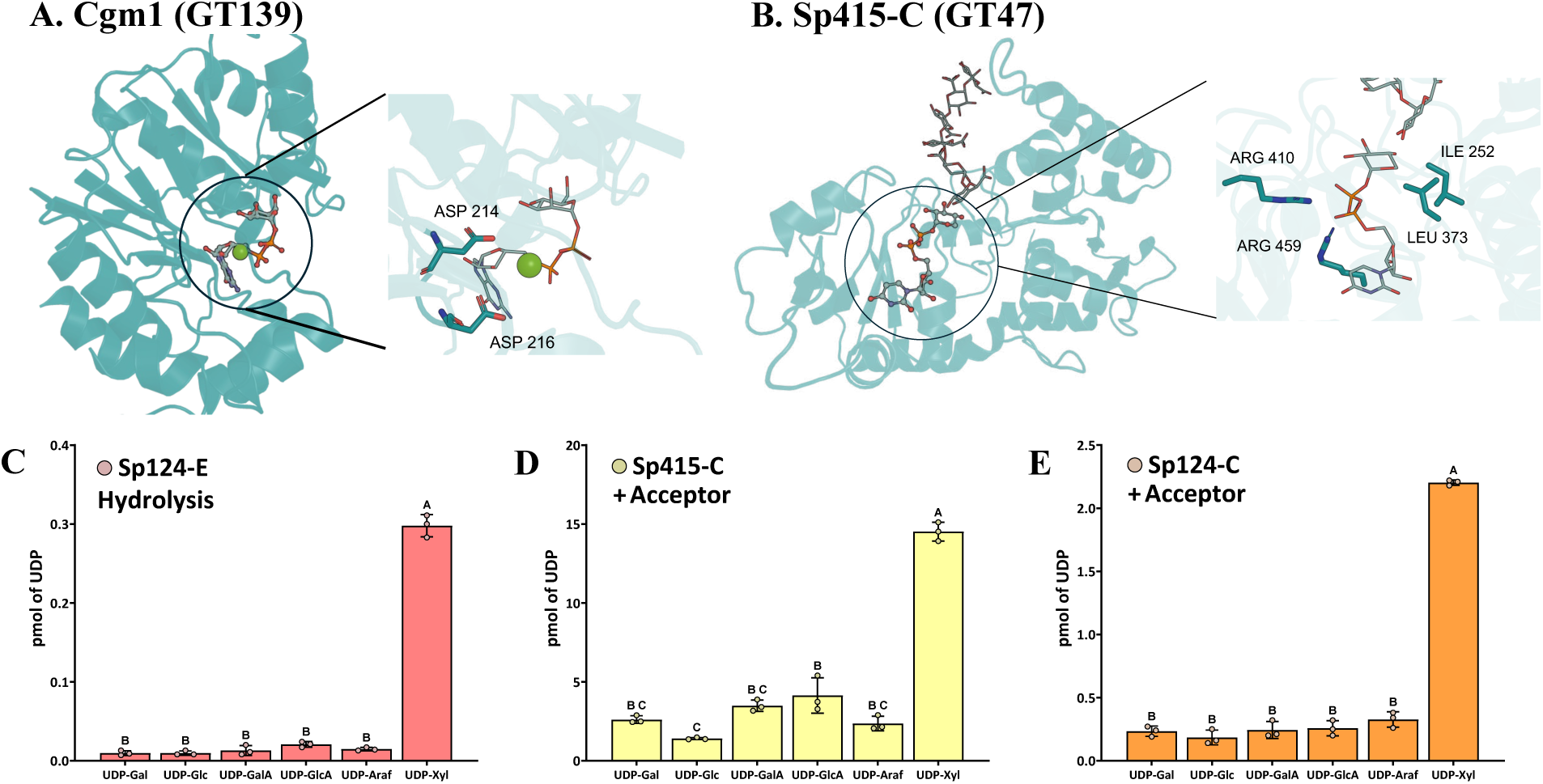
Structural models and experimental validation of donor prediction in uncharacterized plant GTs. **A.** Model of Cgm1 (GT 139) docked with the predicted donor GDP-mannose and manganese, generated by AlphaFold3 (left), and active-site structure showing the DXD motif interacting with the diphosphate group (right) **B.** Model of Sp415-C (GT 47) docked with the predicted donor UDP-xylose and its acceptor homogalacturonan (DP 6), generated by AlphaFold3 (left), and active-site structure showing the two arginines interacting with the diphosphate (right). **C, D, E.** Glycosyltransferase activity in the absence (C) or presence of homogalacturonan oilgosaccharides (DP 5-15) as acceptors in an overnight reaction Sp415-C (**D**) and Sp124-C (**E**). Data is represented as means ± standard deviation of three biological replicates. Individual data points are shown. Letters indicate statistical significance accomplished by a one-way ANOVA followed by a Tukey’s multiple comparison test (*P*≤0.001).

For those with GT-B folds, Glydentify correctly identified UDP-Xyl as the most probable donor for Sp124-E, Sp415-C, and Sp124-C, with prediction scores ranging from 73.4% to 93.3%, in full agreement with our biochemical assays (**Figure 3C, 3D** and **3E**). Likewise, Sp146-A and Sp197-A, were predicted to use UDP-Gal with high confidence (99%), again in agreement with biochemical evidence where those were characterized to act on UDP-Gal^26^(**Supplementary Figure 1**). We also observed that B3GLCT, a member of the GT31 family, was not correctly predicted despite its high sequence similarity to other GT31 enzymes in the training set (**Supplementary Figure 2**).

### Glydentify leverages structural, evolutionary, and biochemical cues for donor recognition

To assess whether Glydentify learns generalizable principles of donor recognition, we conducted a series of interpretability analysis inspired by prior works^27^ demonstrating that attention-based models can recover latent structural determinants of enzyme function. In most enzymes, substrate specificity arises from a complex interplay of interactions: first-shell contacts that form direct hydrogen bonds, ionic interactions, or steric complementarity with the ligand; second-shell residues that do not contact the substrate directly^28^ and even long-range or allosteric effects that influence active-site dynamics through cooperative networks across the protein scaffold^29^. These multi-layered interactions collectively shape substrate binding and catalysis, and often cannot be inferred from primary sequence alone. We therefore examined whether Glydentify, trained solely on sequence embeddings, nevertheless recapitulates these canonical principles of enzyme–substrate recognition in its internal representation of donor-sugar specificity.

We analyzed the attention scores derived from the cross-attention module to identify which residues drive donor-specific recognition. From each selected sample, we extracted the per-residue attention weights 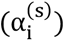 from the output of the cross-attention layer, where *i* the indices correspond to residues in the aligned sequence. To investigate what signals Glydentify relies on to infer donor sugar specificity, we mapped the residue-level attention scores onto predicted three-dimensional structures and analyzed their spatial distribution.

We first performed a global analysis across true-positive GT–donor pairs by plotting the spatial distribution of high-attention residues (normalized attention score > 0.8) relative to the donor molecule. Interestingly, for the GT-A fold, attention was concentrated in residues located primarily within ∼10–25 Å (peak at 16.6 Å) of the donor (**Figure 4A**). Given that distances are measured from the backbone Cα, this indicates that Glydentify prioritizes the immediate active site environment and the surrounding structural scaffold. While for the GT-B fold, attention scores exhibited a bimodal distribution (peaks at 13.6 Å and 22.7 Å) (**Figure 4B**). The proximal peak (∼13.6 Å) highlights residues surrounding the active sites, while the distal peak (∼22.7 Å) likely captures the flexible hinges that allow the enzyme to close around and trap the donor. Detailed per-donor attention concentration distributions showing these consistent fold-specific trends can be found in **Supplementary Figure 3**. Additionally, we found instances where Glydentify uses the biochemical cue to predict the correct glycosyltransferase-donor sugar pair. For example, in glucuronosyltransferases (CAZy family GT 43), residues with high-attention score were found proximal to the negatively charged, C5-carboxylate of UDP-GlcA, highlighting basic side chains positioned within 4-6 Å from carboxylate (**Figure 4C**). This ionic interaction between residues and donor sugar, captured by attention score pairs, suggests that Glydentify understands the biochemical cues of donor sugar recognition by GTs. Together, these structural and quantitative analyses indicate that the model emphasizes residues in spatial proximity to the donor, capturing biochemical cues relevant to sugar specificity.

**Figure 4:**
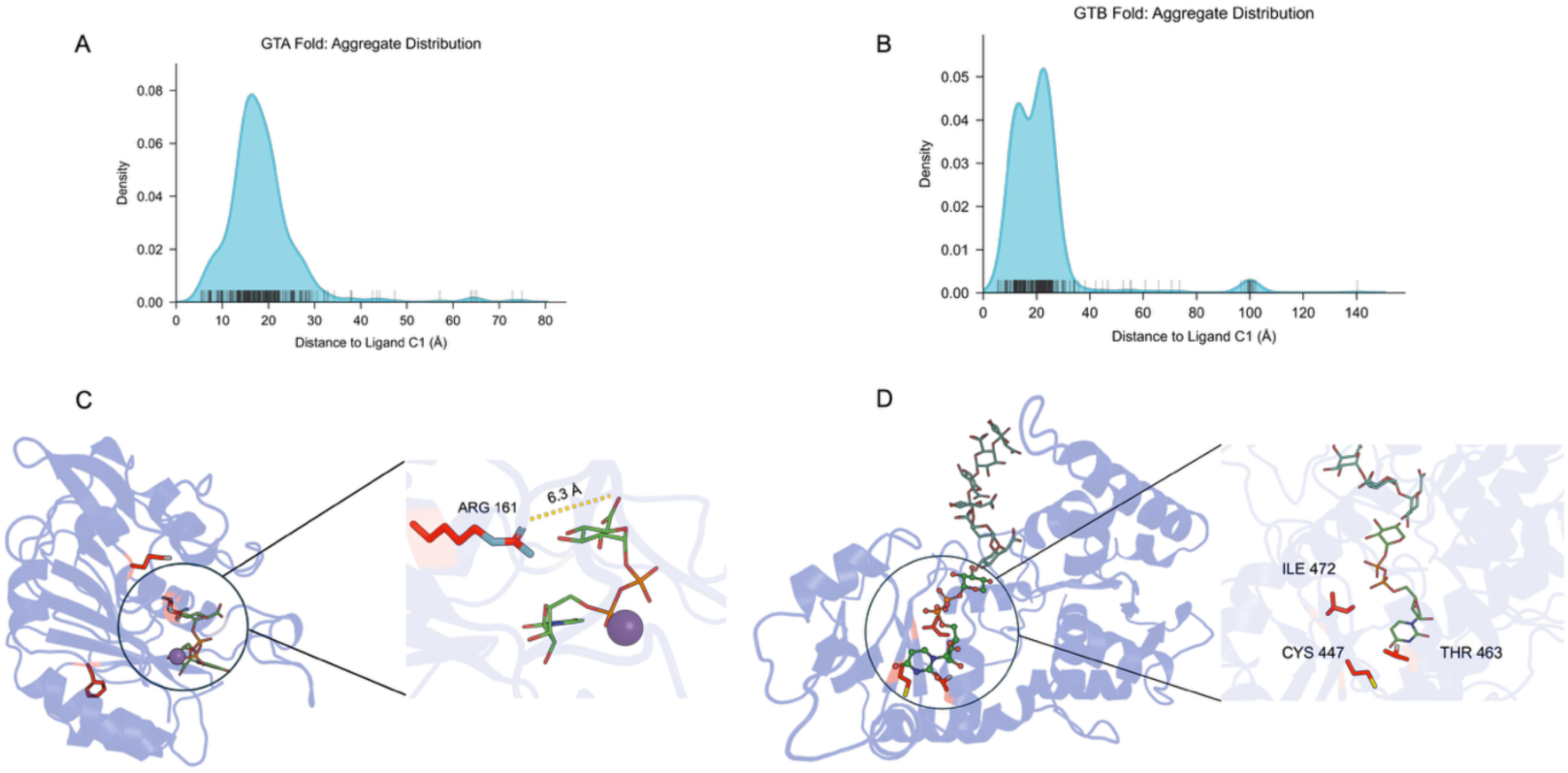
Explainability analysis of donor prediction using attention. **A, B.** Structural colocalization of model attention with the donor binding site. Kernel density estimates show the distribution of Euclidean distances between high-attention residues (normalized score > 0.8) and the donor substrate for (**A**) GT-A and (**B**) GT-B fold enzymes. Analysis was performed on true positive samples using AlphaFold 3-generated structures. Distances were measured from the residue α-carbon (Cα) to the ligand anomeric carbon (C1). **C.** AlphaFold3–predicted structure of B3GAT3 (Uniprot ID: Q9WU47) modeled with UDP-glucuronic acid and Mn²⁺ reveals that an arginine, the residue with the highest cross-attention score, positioned adjacent to the C5-carboxylate of UDP-GlcA. The proximity of this basic residue to the acidic C5-carboxylate suggests a strong ionic interaction is the main determinant of GlcA specificity. **D.** AlphaFold3-predicted ternary complex of a representative GT-B GT47 enzyme (Sp415-C) with UDP-Xyl and acceptor galacturonan mapped with Glydentify attention scores. High-attention residues across GT47 enzymes were found at spatially aligned positions, suggesting that these conserved sites co-evolved to shape donor-sugar specificity.

The GT-47 family exemplified that donor sugar specificity can be shaped by more complex principles rather than by first-shell interactions alone (**Figure 4D**). Instead of focusing on residues in direct contact with the donor sugar, Glydentify consistently highlighted a set of spatially aligned positions outside the donor sugar binding pocket, suggesting that Glydentify may detect co-evolutionary signals rather than simple contact-based features (**Supplementary Figure 4 and supplementary Table 3**). A striking example is a conserved cysteine located in a highly variable region of the central beta-sheet strands, present in all GT-47 homologs, forming disulfide bonds (**Supplementary Figure 5** and **6**) with a spatially proximal cysteine. In Galactosyltransferase Sp146-A and Sp 197-A, a spatially aligned phenylalanine residue is consistently found in the vicinity of this conserved cysteine, suggesting a co-evolved, second-shell pair interaction for recognizing UDP-Gal. Conversely, the first spatially aligned position is occupied by hydroxyl amino acids, serine and threonine, accompanied by branched hydrophobic residues (Ile and Val) at the second spatially aligned position, again adjacent to the conserved cysteine (**Supplmentary Figure 6**). Together, these observations suggest that donor specificity in the GT-47 family may be a result of co-evolved second-shell residue networks.

Overall, our analysis of residues with high attention scores revealed that Glydentify does not rely on a single simplistic feature; instead, the model integrates multiple layers of information, including direct biochemical interactions, structural context, and patterns conserved across evolution to predict donor specificity.

## Discussion

Our study introduces Glydentify, a deep learning framework for predicting glycosyltransferase (GT) donor specificity. By training directly on noisy UniProt annotations and evaluating against manually curated CAZy entries, we show that reliable donor-sequence relationships can be learned from large, imperfect datasets. Previous ML-based approaches have been restricted to specific folds or families, such as the GT-A^10^ or GT-1 family^9,11^, limiting their scalability. General enzyme–substrate predictors such as ESP, EZsepecificity^16,17^ achieve strong results across diverse reaction classes but underperform for GTs. Our models achieve PR-AUCs of 0.8638 for GT-A and 0.9124 for GT-B, surpassing general enzyme–substrate predictors and performing on par with, or better than, curated family-specific approaches. To our knowledge, Glyidentify is the first end-to-end model designed to learn donor usage across the entire sequence space of the GT without requiring extensive manual feature engineering. A major contributor to Glydentify’s performance is the use of SaProt embeddings, which encode 3D structural context absent in sequence-only models. Integrating SaProt representations with UniMol sugar geometries consistently improved accuracy for well-supported donors, demonstrating that the choice of protein language model significantly influences GT donor classification and that structural priors are crucial for distinguishing sugar donors. Furthermore, the model even captured the donor sugar promiscuity of glycosyltransferases (**Supplementary Figure 7** and **8**). For instance, the processive diacylglycerol β-glycosyltransferase from *Mycoplasma genitalium* was predicted to utilize both UDP-Glc (93.5%) and UDP-Gal (99.2%) with high confidence (**Supplementary Figure 7**), consistent with kinetic measurements showing Km = 50 μM for UDP-Glc and Km = 243 μM for UDP-Gal^30^.

Despite the overall strong performance, we observed two notable exceptions in the GT-A model: UDP-GlcNAc and UDP-GalNAc showed negligible or slightly reduced performance gains. This is likely due to a combination of their subtle chemical differences and the limitations of UniProt-derived labels.

GlcNAc and GalNAc differ only at the C4 stereocenter, suggesting that it is difficult for UniMol to separate the epimeric difference. In addition, donor annotations for these sugars in UniProt are often inferred rather than experimentally confirmed, which may further contribute to the lower accuracy observed for these two donors. We also observed that B3GLCT, a member of the GT31 family, was not correctly predicted despite its high sequence similarity to other GT31 enzymes in the training set (**Supplementary Figure 2**), suggesting that high global sequence similarity within GT31 does not always translate into clear donor-specific signals, and closely related sequences with divergent donor sugar specificity may cause difficulty for the model to resolve during training.

UniProt labels contain substantial noise from homology-based inference, and CAZy covers only a small fraction of GT–donor space. Rare donor classes continue to present challenges, with many represented by fewer than ten validated sequences. Our analysis of these minor donors, such as UDP-GalNAc, further confirms this challenge: even with a specialized architecture pre-trained on other diverse sugars, including UDP-GlcNAc, performance remains unattainable with limited data, indicating the strict stereochemical determinants of specificity cannot be robustly migrated from chemically similar donors without sufficient training samples (**Supplementary Figure 10**). With the current depth and quality of available annotations, this likely represents the practical performance ceiling achievable from existing resources. Future advances will depend on the generation of higher-quality, large-scale data—particularly from emerging high-throughput assays capable of directly profiling GT activity. Such datasets could provide the clean, experimentally grounded labels needed for models to learn more accurate sequence–function mappings and iteratively refine themselves through feedback from correct predictions. Combined with improved structural modeling, such as explicit integration of active-site representations, these developments promise to substantially enhance both the accuracy and biological interpretability of GT donor prediction. More broadly, the framework exemplifies how deep learning can close sequence-to-function gaps in enzymology, particularly in cases where data are abundant but noisy, and curated annotations remain scarce.

## Methods

### Data Collection and Preprocessing

We assembled a large-scale training set of GT sequences with donor annotations from UniProt by querying major donor-type enzyme names (e.g. “glucosyltransferase,” “galactosyltransferase,” etc.). To ensure annotation quality, we retained only entries with an annotation score greater than 3, a threshold that reflects substantial manual curation and reduces the prevalence of purely computationally inferred labels. This initial query yielded 126,451 sequences annotated with at least one donor sugar.

We assigned each sequence to the GT-A or GT-B structural fold using hidden Markov model (HMM) profiles from Pfam (PF00535 for GT-A, PF01062 for GT-B) and the HMMER suite (E-value < 1e-5)^31,32^. From the UniProt dataset, we identified 11,492 GT-A and 15,235 GT-B sequences. To train our model efficiently and focus on the catalytic core, each candidate training sequence was cropped to the detected HMM domain, plus 15 flanking residues at both the N- and C-termini of the conserved catalytic domain.

Because donor classes were highly imbalanced, we reduced sequence redundancy within each donor type using mmseqs^33^ clustering at identity thresholds tailored to class prevalence: 95% for donor types with > 500 sequences and 100% for <100 sequences. After selecting one representative per cluster, the final training set comprised 4,938 GT-A and 7,611 GT-B sequences.

For evaluation, we built a high-confidence test set from CAZy by scraping all experimentally characterized GT entries^34^. Since CAZy does not include explicit donor sugar annotations, we utilize a Large Language Model (LLM) to infer the donor sugar used from the protein name or the related publication abstract. GPT 5.1 was used with temperature set to 1.0. The system prompt of the LLM included the input specifications and some examples of inputs and expected output pairs. In case an abstract is provided, the LLM was instructed to extract a subset of lines as evidence showing where the donor sugars are mentioned. The full prompt and code are provided in the **supplementary data**. We clustered sequences at 90% identity and then removed any remaining training sequences with an identity greater than 90% to a test sequence to prevent homology leakage. The resulting test set contains 174 GT-A and 194 GT-B sequences. All test sequences were retained at full length to evaluate the model’s performance on complete proteins without additional truncation.

To further enrich our representations with three-dimensional context, we incorporated predicted structures for all GT sequences. Because the vast majority of these proteins lack experimentally resolved coordinates in the Protein Data Bank, we generated models using AlphaFold^20^. Predictions were performed with default parameters, and the resulting coordinate files served as the structural inputs for downstream representation learning and model evaluation.

To obtain 3D molecular inputs for the donor encoder, we extracted the exact donor sugar molecule from the UniProt “catalytic activity” or “reaction” fields when available; if no explicit annotation existed, we assigned the most common sugar for that enzyme class (e.g. “glucosyltransferase” → UDP-Glucose).

Each SMILES string was converted to an energy-minimized 3D conformer using RDKit, then encoded by UniMolV2 to produce a fixed donor embedding.

Because we only observe positive GT-donor annotations in UniProt and CAZy, we treat every unannotated GT-donor pair as a negative example during training. Concretely, if a sequence is labeled with *k* donors out of the *C* total in our vocabulary (*C* ≈ 10), we generate (*C* − *k*) negatives by pairing that sequence with each of the other donors.

We acknowledge that some true GT–donor pairs may be missing from UniProt/CAZy annotations, resulting in them being erroneously labeled as negatives. However, these false negatives are likely rare for our focused set of major donors. UniProt and CAZy both aim for high coverage of well-studied GTs and any remaining noise is mitigated by the large number of true negatives. In practice, this simple closed-world assumption yielded stable training and reliable generalization to the high-confidence CAZy test set.

### Encoding GTs and Donor Sugars

Given a protein of length *L*, we tokenize the input as:

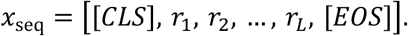

The SaProt encoder produces contextual embeddings

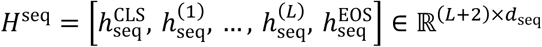

where *h* denotes encoder outputs. We use the [*CLS*] embedding as a global sequence summary and the residue embeddings 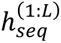 for token–leve interaction. Similarly, for each donor *d* ∈ {1, …, *C*}, we obtain an RDKit conformer from its SMILES and feed it to UniMol V2 to obtain atom–level embeddings.

UniMol uses special tokens as well:

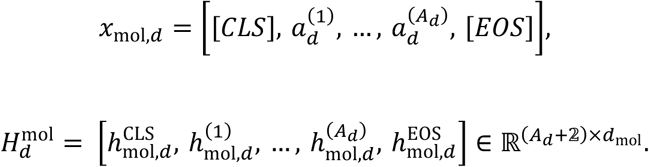

Both Saprot and UniMol as well as the generated embeddings are frozen to reduce overfitting and training cost. To adapt embeddings, map raw encoder embeddings *h* to adapter outputs *z* in their native dimensions via:

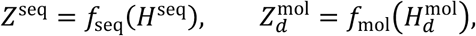

where *f*_seq_ *and f*_mol_ are MLP (Multi-Layer Perceptrons) layers consist of a linear projection followed by a GELU activation and Layer Normalization. We write:

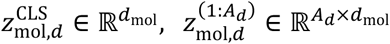

### Bi-directional cross-attention module and classification head

We compute donor–specific interactions in both directions. In *mol* → *seq*, the donor [*CLS*] queries residue tokens; in *seq* → *mol*, the protein [*CLS*] queries atom tokens. Importantly, keys/values exclude [*CLS*] [*EOS*] and use only 1: *L* residues or 1: *A*_2_ atoms:

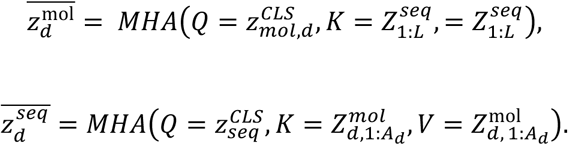

We apply residual connections and layer normalization:

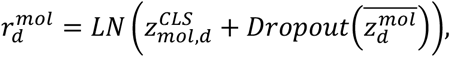

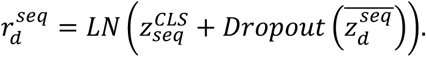

We project each side to a shared width *d*_*_ and fuse:

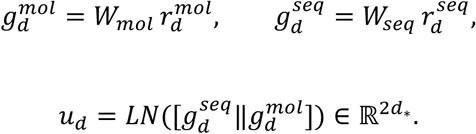

A two–layer MLP produces a donor–specific logit *s*_2_ ∈ *R* and probability 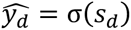

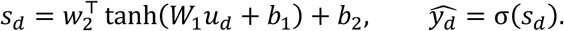

Stacking all donors yields a multi–label prediction vector 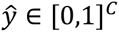.

### Loss function and optimization

Because negative labels vastly outnumber positives, we adopt ASL (Asymmetric Loss)^35^ to down–weight negatives while preserving the positive signal. Let *x* ∈ *R^C^* be the logits, *y* ∈ {0,1}*^C^* the targets, and *p* = σ(*x*). Denote class index by *c* and choose focusing parameters γ_+_ ≥ 0 (positives) and γ_-_ > 0 (negatives). Define an asymmetric clipping margin *m* ∈ [0,1) applied only to the negative term:

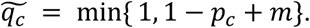

The per–example ASL is:

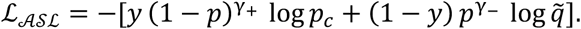

The positive term receives standard or mildly focused weighting via 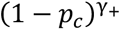, while the negative term is aggressively damped by 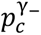 and the clipping 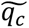, which reduces gradients from easy negatives (small *p_c_*).

### Training Details of Glydentify

We formulated the prediction task as a binary classification problem, where the model takes a glycosyltransferase (GT) sequence and a donor substrate as input pairs. The model was trained using ASL with γ_+_ = 0 and γ_-_ = 2. We optimized the network using AdamW with default parameters, a learning rate of 5×10 −5, and a weight decay of 1×10 −4. The training batch size was set to 16 and ran for a maximum of 30 epochs, utilizing early stopping with a patience of 5 epochs to prevent overfitting.

### Protein Expression and Purification

The expression constructs for all plant GTs used in this study (details in **Table 1**) were generated as truncated catalytic domains as NH2-terminal fusion proteins in the pGEM2 expression vector, as described in prior studie^36^. Briefly, the fusion protein coding region was comprised of a 25-amino acid signal sequence, an His8 tag, AviTag, the “superfolder” GFP, the 7-amino acid recognition sequence of the tobacco etch virus (TEV) protease followed by the truncated coding region of each individual plant GTs under study. Coding sequences for *Spirodela polyrhiza* (v2) and *Spirodela intermedia* were obtained from Phytozome 14 and the National Center for Biotechnology Information, respectively. The truncated catalytic domains of Sp124-E (Spipo24G0012400; amino acids 37-801), Sp415-C (Spipo8G0041500, amino acids 38-564), Sp124-C (Spipo2G0124300, amino acids 25-403), Sp291-D (Spipo1G0029100, amino acids 45-531), and Sl728-C (SICAA7408728.1; amino acids 44-555) without the transmembrane domains (predicted using DeepTMHMM - 1.035) were cloned into pTwist Gateway ENTR via gene synthesis (Twist Biosciences). The genes were transferred into the mammalian expression vector pGEn2-DEST^36,37^ using Gateway^TM^ recombination.

The expression constructs were then used for transient transfection of suspension culture HEK293-F cells (FreeStyleTM 293-Fcells, Thermo Fisher Scientific, Waltham MA) maintained at 0.5–3.0x10^6^ cells/ml in a humidified CO2 platform shaker incubator at 37 °C with 50 % relative humidity and 125 RPM. Transient transfection was performed using HEK293-F cells (2.5-3.0x10^6^ cells/ml) in the expression medium comprised of a 9:1 ratio of FreestyleTM293 expression medium (Thermo Fisher Scientific, Waltham MA) and EX-Cell expression medium including Glutmax (Sigma-Aldrich). Transfection was initiated by the addition of plasmid DNA and polyethyleneimine as transfection reagent (linear 25-kDa polyethyleneimine, Polysciences, Inc., Warrington, PA). Twenty-four hours post-transfection, cell cultures were diluted with an equal volume of fresh media supplemented with valproic acid (2.2 mM final concentration), and protein production was continued for an additional 5 days at 37 °C, 125 RPM and 5% CO2. The cell cultures were harvested, clarified by sequential centrifugation at 1200 rpm for 10 min and 3500 RPM for 15 min at 4 °C, and passed through a 0.8 µM filter (Millipore, Billerica, MA). The crude protein was then subjected for further purification using immobilized metal affinity chromotography (IMAC) using an AKTA 25L system (Cytiva) at 4 °C with HisTrap FF prepacked 5 ml or 1ml columns (Cytiva)^36,37^. Purified proteins were also loaded onto an SDS-PAGE gel to confirm the expected molecular weight. All proteins were buffer exchanged into 50 mM HEPES Sodium Salt (pH 8) with 100 mM sodium chloride and transferred to -80 °C for long-term storage.

### Glycosyltransferase Activity Assay

UDP-sugar donors used in reactions were CIAP-treated (Calf Intestinal Alkaline Phosphatase) according to Sheikh and Wells, 2006^38^ The UDP-Glo™ Glycosyltransferase Assay (Promega) was carried out in a total volume of 10 μL, consisting of 250 mM Tris+MES+MOPS pH 7 or pH 5.5, 0.5 mM UDP-sugar donor, with and without 0.1 mg/mL of acceptor (when indicated), and 1 μM GT enzyme. Reactions were set at room temperature for either 2 hours or overnight as indicated in the legend. For UDP quantitation, 5 μL of the reaction was subsequently mixed with 5 μL of UDP-Glo™ Detection Reagent directly in white polystyrene, low-volume, 384-well assay plates (Corning) and incubated for 60 minutes. Luminescence was recorded with a multifunctional microplate reader BioTek Synergy LX Multi-Mode Reader (Agilent). Data was quantified relative to a standard curve of UDP.

## Data Availability

The raw protein sequences were retrieved from the UniProt database (https://www.uniprot.org). The raw information used to generated testing set can be found in CAZy database (https://www.cazy.org/GlycosylTransferase-family).

## Code Availability

The source code for Glydentify, including model architecture, training scripts, and inference pipelines, as well as the processed training and testing data are is available for peer review at https://github.com/RuiliF/Glydentify. A permanent DOI will be minted via Zenodo upon acceptance.

## Acknowledgements (optional)

This work was supported by funding from the NSF BioFoundry (2400220/U.S. National Science Foundation BioFoundry: Glycoscience Research, Education and Training). Funding for BU from U.S. DOE, Office of Science, BER program, GSP grants DE-SC0023223 and DE-SC0026057 is acknowledged.

## Ethics declarations

### Competing interests

Submission of a competing interests statement is required for all content of the journal.

## Notes

### Competing Interest Statement

The authors have declared no competing interest.

