## Supplementary Files for "Glydentify: An explainable deep learning platform for glycosyltransferase donor substrate prediction"

### Supplementary Text

**Supplementary Text 1.** Description of data efficiency experiment.

Existing glycosyltransferase databases are inherently biased toward common sugar donors. However, identifying enzymes that utilize these rare specificities is a critical bottleneck for expanding the biocatalytic toolbox, underscoring the need for models capable of robust generalization from minimal data. Notably, we observed that model performance for individual donors was not strictly correlated with the number of available training samples. To quantify the amount of data required for generalization to under-represented donors, we designed a systematic data efficiency experiment.

We simulated this scenario by holding out all samples of a specific target donor $d$ from the main training set, noted as $D_{target}^{train}$. We first trained a baseline model (denoted $M_0$) on the remaining data ($D_{base}$) using the same hyperparameter settings as the main models. We then evaluated data efficiency by freezing the encoder and fine-tuning only the projection and classifier layers of $M_0$ on the held-out donor. For a given donor $d$, we constructed training sets containing $k$ positive examples randomly sampled from $D_{target}$ and a corresponding set of negatives sampled from $D_{base}^{train}$, maintaining a 1:5 ratio of positives to negatives. We varied support size $k \in \{0, 1, 5, 10, 20, 50, 100, 200, 500\}$ to cover the spectrum from zero-shot to data-rich settings. For each $k > 0$, we performed 5 independent replicates. The final models were evaluated on the held-out test set of donor $d$ $D_{target}^{test}$ using PR-AUC to identify the saturation point where additional labeling yields diminishing returns. The performance curves for each donor sugar are shown in **Supplementary Figure 10**.

#### Supplementary Figures

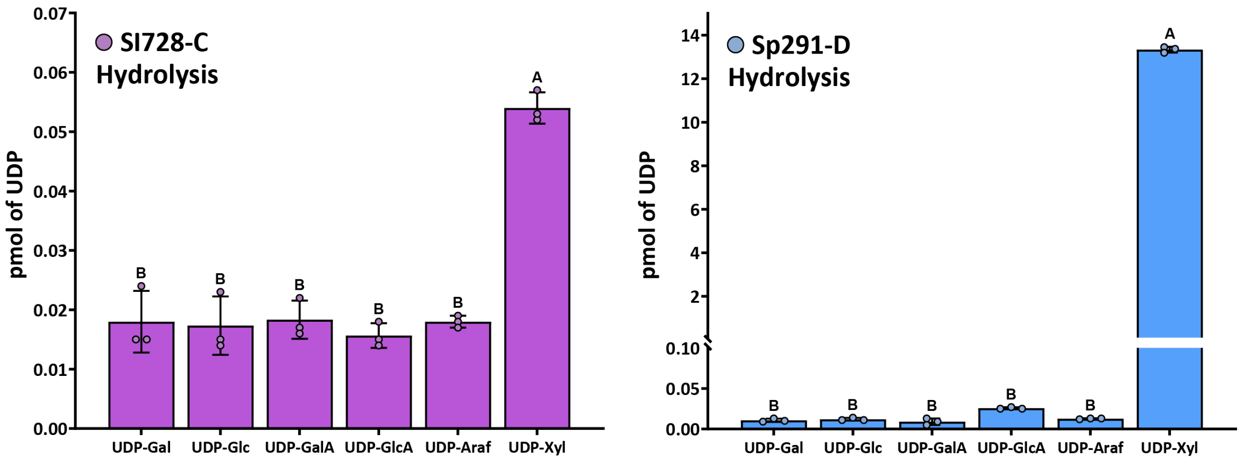


**Supplementary Figure 1. NDP-sugar screen of uncharacterized *S. polyrhiza* GT47 enzymes.** The relative activity of Sl728-C (left, purple) and Sp291-D (right, blue) in the presence of various UDP- sugar donors in the absence of acceptor substrates. Enzymatic activity was measured by quantifying the UDP by-product after enzymatic transfer of a UDP-sugar donor to water (hydrolysis). Error bars indicate Mean ± SD with an n of 3 biological replicates. Letters above indicate statistical significance accomplished by a one-way ANOVA followed by a Tukey's multiple comparison test ($P \leq 0.001$).

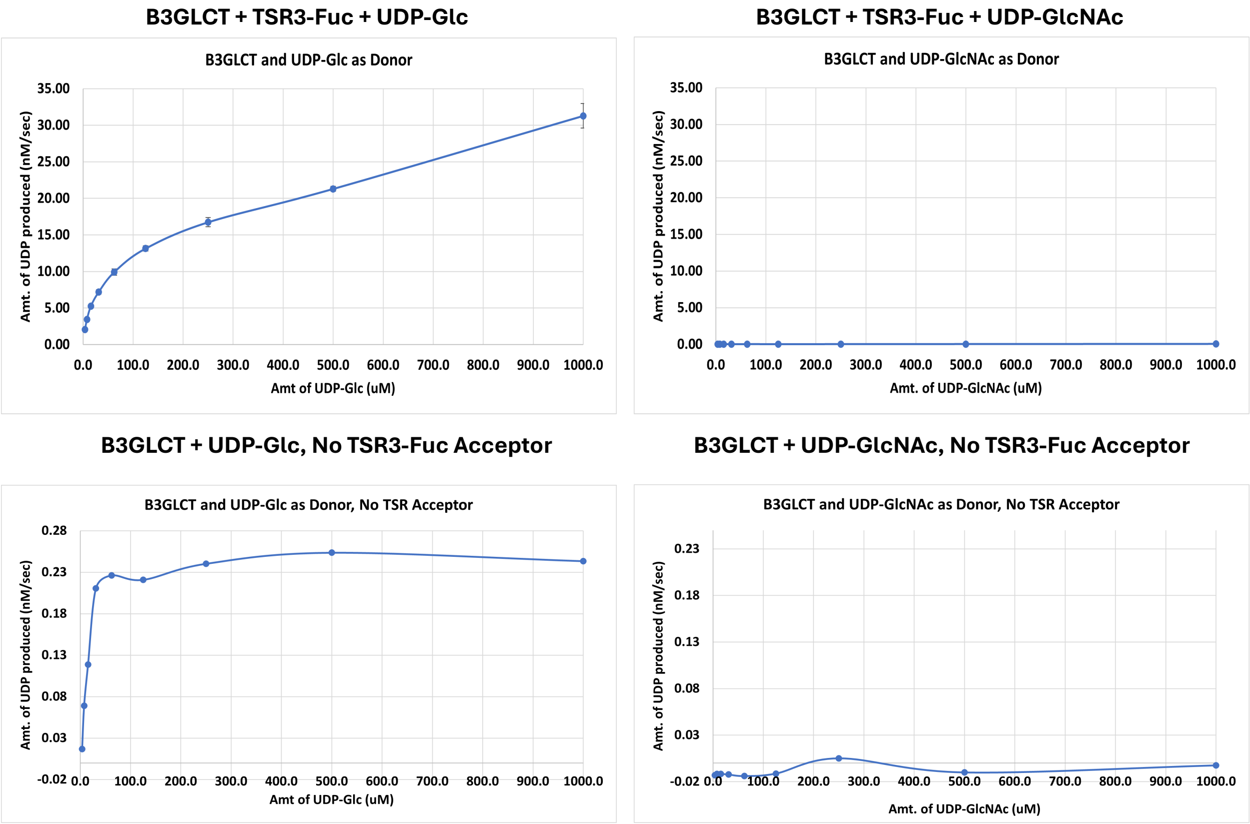


**Supplementary Figure 2. UDP-Glo™–based donor substrate specificity analysis of B3GlcT using UDP-Glc and UDP-GlcNAc.** Reactions were performed with UDP-Glc in the presence of the acceptor TSR3-Fuc (top left) and in the absence of acceptor (bottom left). B3GlcT showed clear activity toward UDP-Glc under both conditions. In contrast, no detectable activity was observed with UDP-GlcNAc, either in the presence of TSR3-Fuc (top right) or in the absence of acceptor (bottom right).

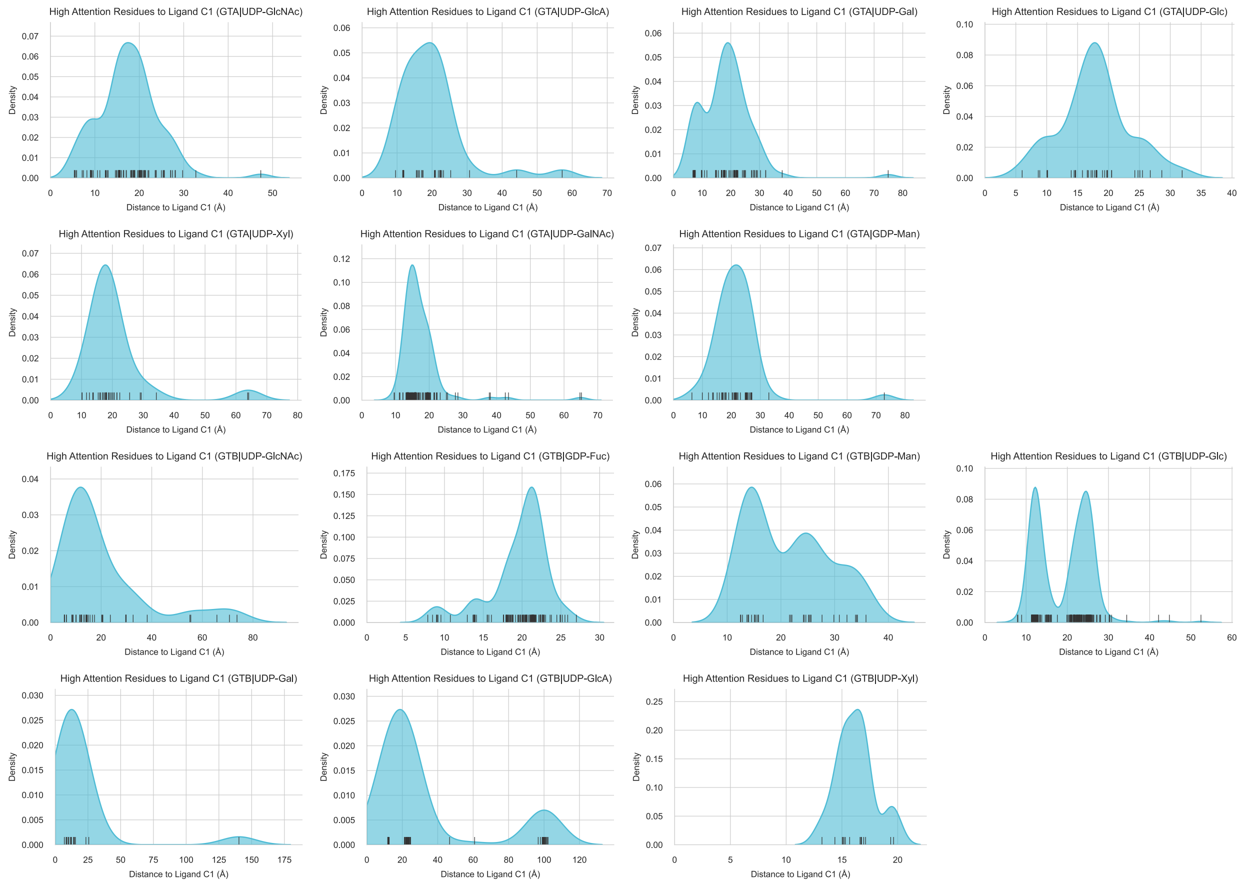


**Supplementary Figure 3. Spatial proximity of high-attention residues to the donor ligand in AlphaFold 3-predicted structures.** Distributions of Euclidean distances (Å) between residues identified as "high attention" by the model and the donor sugar substrate. Analysis is restricted to true positive predictions to validate model interpretability. Distances were calculated between the anomeric carbon (C1) of the donor ligand and the alpha-carbon (Cα) of residues exhibiting a normalized attention score > 0.8 in AlphaFold 3 (AF3) generated structures. Rug plots (black ticks) along the x-axis represent individual residue data points.

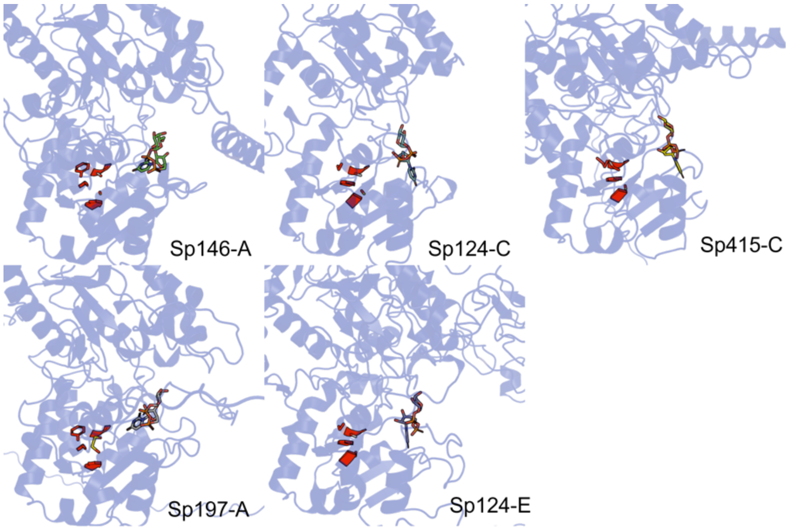


**Supplementary Figure 4. Spatial alignment of high-attention residues across GT-47 enzymes.** Predicted structures of representative GT-47 enzymes showing the top attention-weighted residues (red) mapped onto AlphaFold3 models docked with the predicted donor sugar. Across all enzymes, high-attention residues occupy the same 3D structural positions.

```
Spipo1G0014600|Spipo1G0014600      H-SPSSIMQMFQSSLFCLQPQGDSYTRRSAFDSMLAGCIPVFFHPGSAYTQYTWHLPRNY    419
Spipo17G0019700|Spipo17G0019700    SNRTAETLNLFLGSVFCLQPRGDSFTRRSTFDCMVAGAIPVFFWRRSAYMQYEWFLPPDG    452
Spipo24G0012400|Spipo24G0012400    PRRASDYYSELSSSVFCGVFPGD-GWSGRMEDSILQGCIPVVIQDG---IFLPYENVFNY    681
Spipo3G0015400|Spipo3G0015400      ---IRRAGGGMRASKFCLNIAGDTPSSNRLFDAIASHCVPVIVSDE---IELPYEDVLDY    378
Spipo2G0124300|Spipo2G0124300      ATGELVYQKRFYRTKFCICPGGSQVNSARIADSIHYACVPVVLSDY---YDLPFNDILDW    334
Spipo8G0041500|Spipo8G0041500      -----TYAEHMKSSRYCLCPRGFEVNSPRLAETFFYECVPVIISDN---FVPPFFDVLNW    487
SICAA7408728                       QNATMSYAEYMRSSRYCICPRGYEVHSPRVVEAIFYQCVPAIISDN---FVPPLFEVLNW    445
Spipo13G0013200|Spipo13G0013200    KLGRIDYFHHLRNAKFCLAPRGESSWTLRFYEAFFMECVPVILSDQ---VELPFQNIIDY    361
Spipo1G0029100|Spipo1G0029100      RKRHDGFRSEMARSVFCLCPRGWAPWSPRLVESVAVGCVPVIIADG---IRLPFSDTVRW    362
                                             :  : :*    *         : .   .:*...
```

**Supplementary Figure 5. Multiple sequence alignment of GT-47 enzymes showing a family-wide conserved cysteine at a structurally aligned position.** *Spirodela intermedia* GT47 sequences were subject to Clustal Omega multiple sequence alignment. Family-wide conserved cysteine (highlighted with red box) corresponds to one of the high-attention positions identified by Glydentify. This cysteine forms di-sulfide bond with the other conserved cysteine (highlighted with blue box).

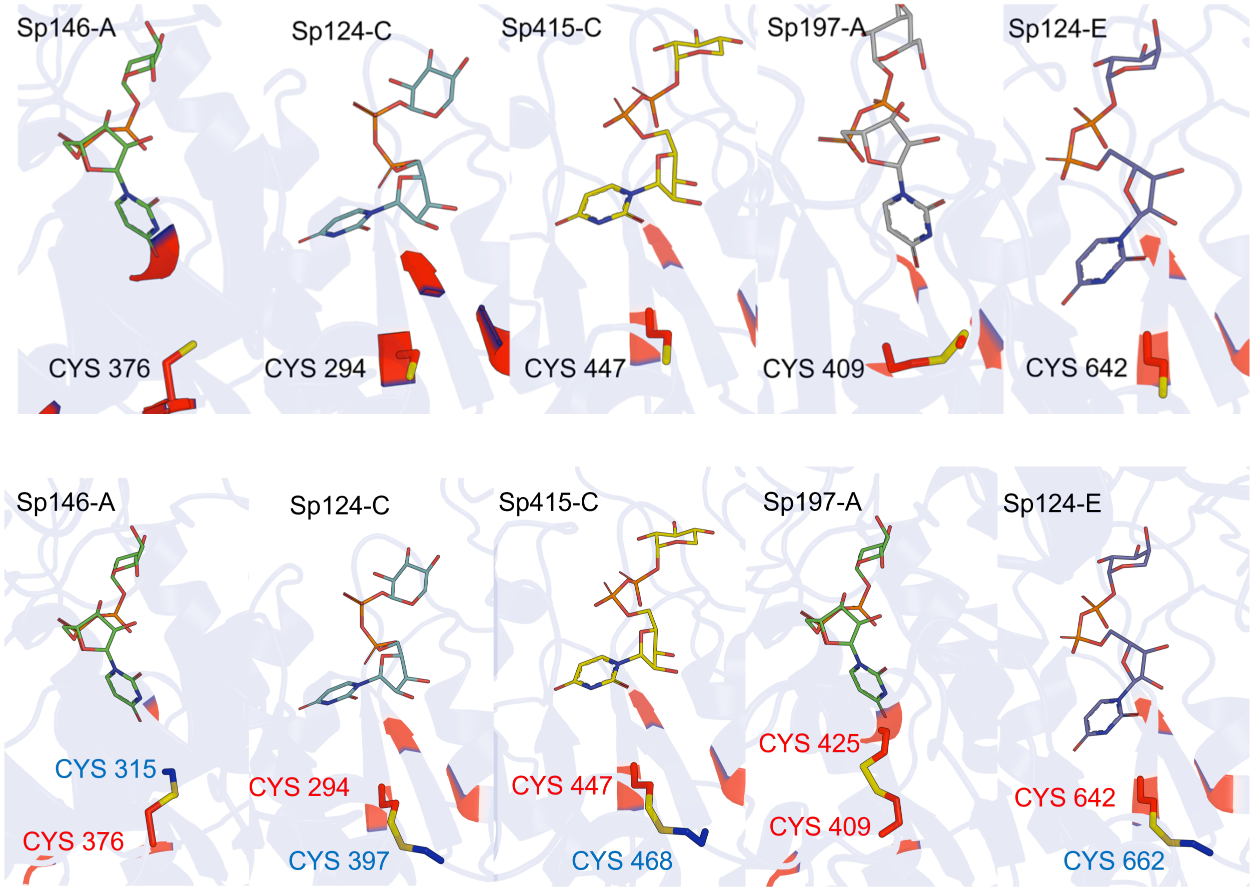


**Supplementary Figure 6. Spatially aligned cysteine across GT-47 enzymes with high-attention score.** Predicted structures of representative GT-47 glycosyltransferases highlighting the family-wide conserved cysteine (red) at a shared 3D structural position are shown. The conserved cysteine is located on the same β-strand of the central Rossmann-like domain in every homolog, indicating that the model captures a family-specific evolutionary constraint.

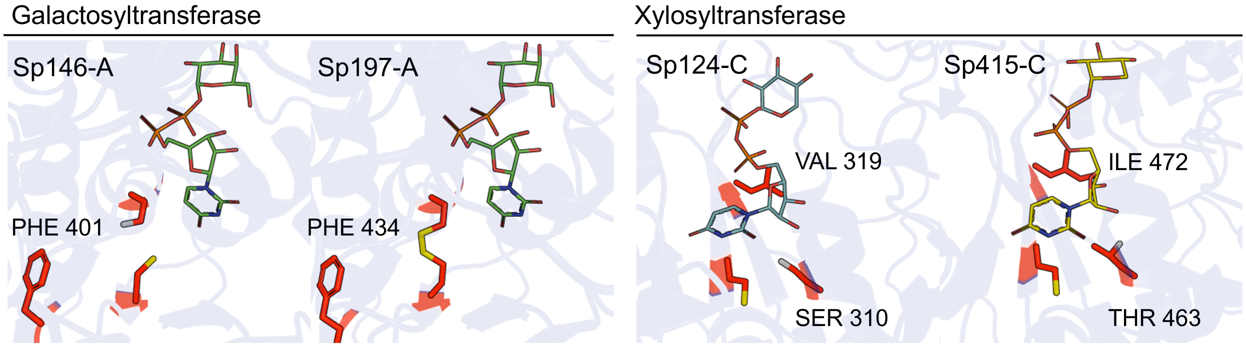


| GT | Residue | Normalized Attention Score |
|---|---|---|
| Sp146-A | PHE 401 | 1.00 |
| Sp197-A | PHE 434 | 1.00 |
| Sp124-C | SER 310 | 0.99 |
| | VAL 319 | 1.00 |
| Sp415-C | ILE 472 | 0.93 |
| | THR 463 | 0.99 |

**Supplementary Figure 7. Spatially aligned residues that may have co-evolved to distinguish sugar donor specificity between GalTs and XylTs.** Representative GT-47 AlphaFold3 structures docked with their predicted sugar donors reveal that galactosyltransferases (left) and xylosyltransferases (right) share a set of spatially aligned positions that differ in residue identity. In GalTs, Phe is consistently found adjacent to the conserved Cys, suggesting a possible co-evolved pair associated with UDP-Gal specificity. In XylTs, Ser/Thr residues, together with Val/Ile at the second aligned position, appear in the same 3D locations, indicating a distinct co-evolutionary pattern linked to UDP-Xyl specificity.

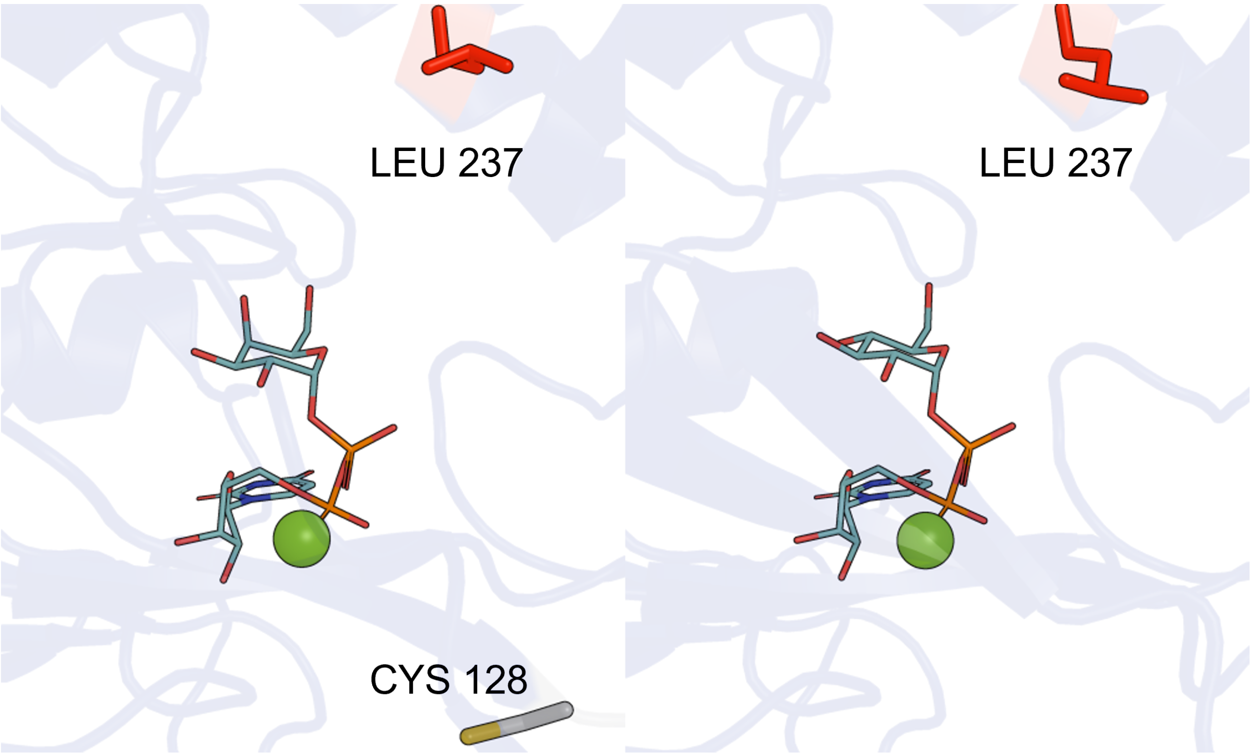


**Supplementary Figure 8. Alphafold3-predicted structure of diacylglycerol β-glycosyltransferase from Mycoplasma genitalium highlighting high-attention residues associated with dual donor-sugar specificity.** AF3 structures of diacylglycerol β-glycosyltransferase (UniProt ID: Q9ZB73) were modeled with UDP-Gal (left) and UDP-Glc (right) docked in the presence of $Mg^{2+}$. Residues assigned high attention by GlyIdentify are identical between the two donor-sugar complexes except for a single cysteine 128 uniquely highlighted in the UDP-Gal model. This pattern illustrates GlyIdentify's ability to capture donor-sugar promiscuity and pinpoint residue-level hotspots associated with promiscuity.

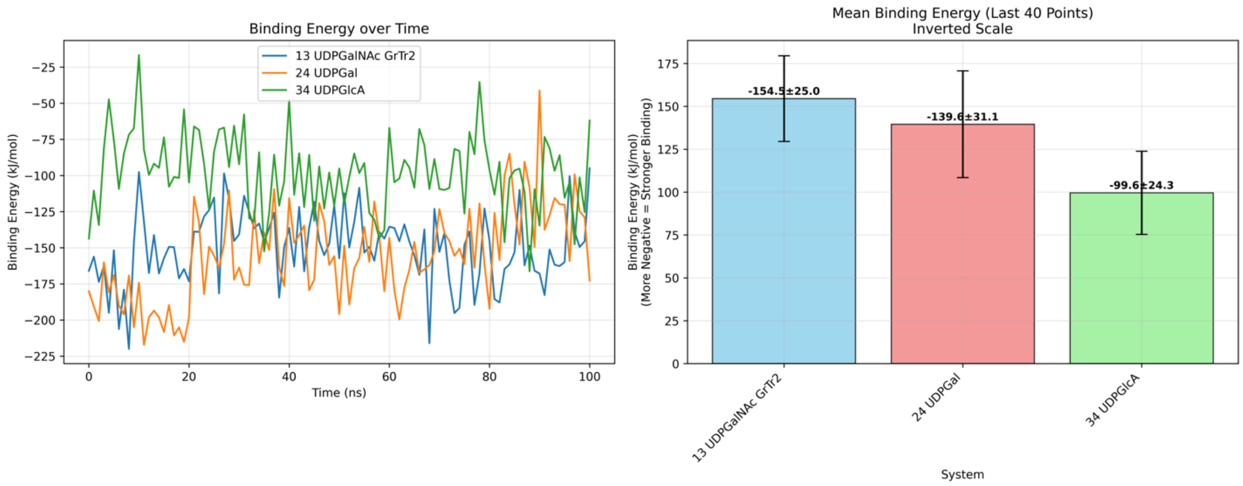


**Supplementary Figure 9. Molecular dynamics simulation of human globoside α1,3-N-acetylgalactosaminyltransferase 1 supports GlyIdentify-predicted dual donor-sugar specificity.** Human globoside α1,3-N-acetylgalactosaminyltransferase 1 (UniProt ID: Q8N5D6) was predicted by GlyIdentify to utilize both UDP-GalNAc and UDP-Gal with high confidence. To evaluate this predicted dual specificity, 100-ns MD simulations were performed for complexes containing UDP-GalNAc, UDP-Gal, or a negative-control donor, UDP-GlcA. Binding-energy analysis (left) and mean binding energies from the final 40-ns equilibrated window (right) show that both UDP-GalNAc and UDP-Gal form substantially more favorable binding interactions than UDP-GlcA. These results support the Glyidentify's prediction on human globoside α1,3-N-acetylgalactosaminyltransferase 1 that may exhibit dual donor-sugar specificity.

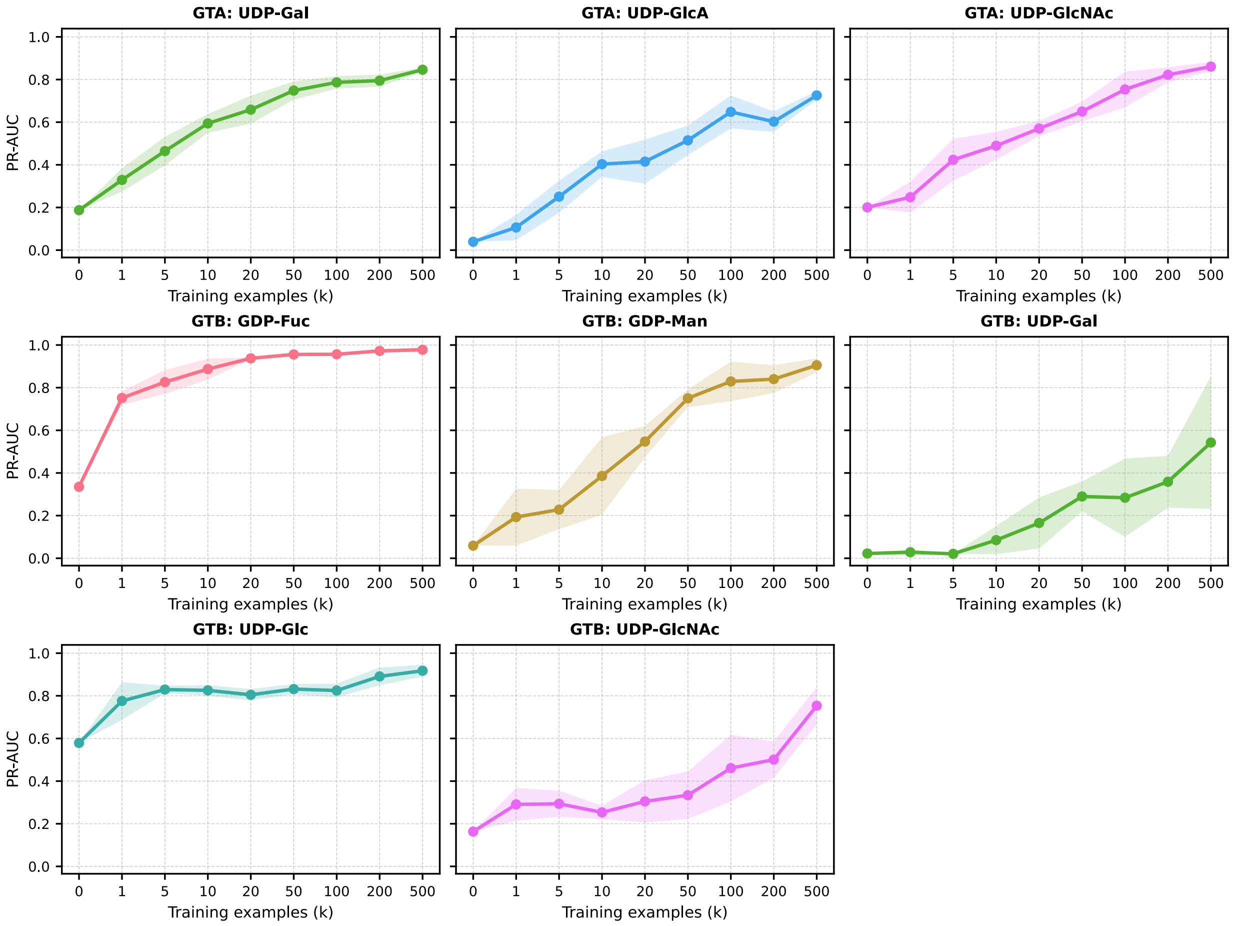


**Supplementary Figure 10. Data efficiency and few-shot adaptation of Glydentify to novel donor substrates.** Performance (PR-AUC) of the Glydentify model after fine-tuning on held-out donor substrates using k training examples. Analysis was restricted to donors with at least 500 positive samples in the original training set. Each curve represents a different held-out donor, with solid lines showing mean PR-AUC across 5 independent replicates and shaded regions indicating standard deviation. The x-axis shows training set size at discrete checkpoints (k ∈ {0, 1, 5, 10, 20, 50, 100, 200, 500}) to emphasize performance in the few-shot learning regime. k = 0 represents the pre-trained model performance without fine-tuning.

### Supplementary Tables

**Supplementary Table 1.** Complete Overall Performance on CAZy dataset.

| *Fold* | *Model* | *ROC-AUC* | *PR-AUC* | *F1 micro* | *MCC* |
|---|---|---|---|---|---|
| *GT–A* | ESP | 0.6509 | 0.1664 | 0.2065 | 0.0932 |
| | EzSpecificity | 0.4189 | 0.1252 | 0.0081 | 0.0597 |
| | ESM2-650M + MLP | 0.7741 | 0.3787 | 0.3333 | 0.2319 |
| | ESM-C 600M + MLP | 0.9230 | 0.6827 | 0.5714 | 0.5253 |
| | SaProt-650M-AF2 + MLP | 0.7565 | 0.2806 | 0.3199 | 0.2160 |
| | Glydentify (ESM2) | *0.9543* | 0.8101 | **0.7928** | **0.7680** |
| | Glydentify (ESM-C) | 0.9468 | *0.8164* | 0.7352 | 0.7002 |
| | Glydentify (SaProt) | **0.9601** | **0.8636** | *0.7863* | *0.7583* |
| *GT–B* | ESP | 0.7459 | 0.2027 | 0.1931 | 0.1630 |
| | EzSpecificity | 0.5797 | 0.1646 | 0.0 | -0.125 |
| | ESM2-650M + MLP | 0.9575 | 0.8183 | 0.7221 | 0.6993 |
| | ESMC-600M + MLP | 0.9638 | 0.8622 | 0.7766 | 0.7549 |
| | SaProt-650M-AF2 + MLP | 0.9119 | 0.7300 | 0.6341 | 0.6027 |
| | Glydentify (ESM2) | *0.9683* | **0.9110** | **0.8562** | **0.8399** |
| | Glydentify (ESMC) | **0.9684** | 0.9042 | 0.8272 | 0.8082 |
| | Glydentify (SaProt) | 0.9636 | *0.9090* | *0.8523* | *0.8357* |

**Supplementary Table 2.** Performance on minor (training sample less than 50) donors.

| *Fold* | *Donor Sugar* | *# Training Samples* | *# Testing Samples* | *# True Positives* | *# False Negatives* |
|---|---|---|---|---|---|
| *GT–A* | UDP-GalA | 27 | 2 | 1 | 1 |
| | UDP-Rha | 6 | 0 | N/A | N/A |
| | dTDP-Rha | 1 | 3 | 0 | 3 |
| *GT–B* | UDP-Rha | 3 | 9 | 0 | 9 |

**Supplementary Table 3.** Interpretability of GT47 predictions.

| NAME | SUGAR DONOR | SPATIALLY ALIGNED POSITION A | SPATIALLY ALIGNED POSITION B | SPATIALLY ALIGNED POSITION C |
|---|---|---|---|---|
| **SP146-A** | UDP-Gal | Ser 392 | PHE 401 | CYS 376 |
| **SP197-A** | UDP-Gal | CYS 425 | PHE 434 | CYS 409 |
| **SP124-E** | UDP-Xyl | SER 657 | VAL 666 | CYS 642 |
| **SP415-C** | UDP-Xyl | Thr 463 | ILE 472 | CYS 447 |
| **SP124-C** | UDP-Xyl | SER 310 | VAL 319 | CYS 294 |
